# Heterologous glycosyl hydrolase expression and cellular reprogramming resembling sucrose-induction enable *Zymomonas mobilis* growth on cellobiose

**DOI:** 10.1101/854646

**Authors:** Nagendra P. Kurumbang, Jessica M. Vera, Alexander S. Hebert, Joshua J. Coon, Robert Landick

## Abstract

Plant derived fuels and chemicals from renewable biomass have significant potential to replace reliance on petroleum and improve global carbon balance. However, plant biomass contains significant fractions of oligosaccharides that are not usable natively by many industrial microorganisms, including *Escherichia coli*, *Saccharomyces cerevisiae,* and *Zymomonas mobilis*. Even after chemical or enzymatic hydrolysis, some carbohydrate remains as non-metabolizable oligosaccharides (*e.g*., cellobiose or longer cellulose-derived oligomers), thus reducing the efficiency of conversion to useful products. To begin to address this problem for *Z. mobilis*, we engineered a strain (*Z. mobilis* GH3) that expresses a glycosyl hydrolase (GH) with β-glucosidase activity from *Caulobacter crescentus* and subjected it to an adaptation in cellobiose medium. Growth on cellobiose was achieved after a prolonged lag phase in cellobiose medium that induced changes in gene expression and cell composition, including increased expression and secretion of GH. These changes were reversible upon growth in glucose-containing medium, meaning they did not result from genetic mutation but could be retained upon transfer of cells to fresh cellobiose medium. After adaptation to cellobiose, our GH-expressing strain was able to convert about 50% of cellobiose to glucose within 24 hours and use it for growth and ethanol production. Alternatively, pre-growth of *Z. mobilis* GH3 in sucrose medium enabled immediate growth on cellobiose. Proteomic analysis of cellobiose- and sucrose-adapted strains revealed upregulation of secretion-, transport-, and outer membrane-related proteins, which may aid secretion or surface display of GHs, entry of cellobiose into the periplasm, or both. Our two key findings are that *Z. mobilis* can be reprogrammed to grow on cellobiose as a sole carbon source and that this reprogramming is related to a natural response *of Z. mobilis* to sucrose that enables sucrose secretion.

## INTRODUCTION

Advances in synthetic biology and lignocellulosic hydrolysate production have encouraged development of the α-proteobacterium *Zymomonas mobilis* as a platform microbe for production of renewable biofuels and chemicals (*e.g*., ethanol, C_4_ and C_5_ alcohols, or C_5_-C_15_ terpenoids) from lignocellulosic biomass (1–3). Efficient conversion of lignocellulose to biofuels and bioproducts is essential for the development of sustainable sources of fuels and chemicals that minimize competition with food production (4). *Z. mobilis* is promising for lignocellulosic conversions because it rapidly and efficiently converts glucose to ethanol, tolerates high ethanol concentrations, tolerates other inhibitors present in the lignocellulosic hydrolysates, and has a small (2.1 Mb) and increasingly well-defined genome amenable to synthetic biology approaches (5). *Z. mobilis* uses the Entner–Doudoroff pathway for glycolysis, which reduces the amount of protein synthesis required to convert glucose to pyruvate relative to Embden–Meyerhof–Parnas glycolysis used by yeast and many other microbes (6). Additionally, fermentation to ethanol is not tightly linked to cell growth, enabling exceptionally high flux that continues even when cell growth stops.

Despite its potential as a platform microbe for lignocellulosic biofuel production, wild-type *Z. mobilis* has a limited substrate range consisting of glucose, fructose, and sucrose, and thus requires engineering to convert the diverse sugar monomers and oligomers present in lignocellulosic hydrolysates. Engineered strains that convert the 5-carbon sugars xylose (7) and arabinose (8) or cellobiose (9) to ethanol have been developed, but use of sugar oligomers other than its native substrate sucrose at levels that support *Z. mobilis* cell growth has not been reported. Sugar oligomers including cello- and xylo-oligomers can be significant components of lignocellulosic hydrolysates, representing about 18-25% of total soluble sugars in corn stover hydrolysates, especially when hydrolysates are generated under milder conditions that minimize inhibitor production or enable recovery of intact lignin (10). Residual glycosyl hydrolases (GHs) in enzymatically prepared hydrolysates may enable conversion of these oligomers to monomers as conversion reduces end-product inhibition of GHs, but there is growing interest in enzyme-free chemical deconstruction methods like those enabled by the renewable solvent γ-valerolactone (11). These enzyme-free hydrolysates will lack the residual GHs necessary for continued oligomer hydrolysis during fermentation. Thus, equipping *Z. mobilis* with an ability to convert sugar oligomers efficiently by production and secretion of GHs would improve prospects for its use as a platform microbe for lignocellulosic conversions.

As cellobiose is the most fundamental unit of these unusable sugar oligomers, we targeted production and secretion of GHs that would enable efficient cellobiose conversion by *Z. mobilis* by heterologous expression of *Cellvibrio japonicus* Cel3A and *Caulobacter crescentus* CC_0968, both belonging to glycosyl hydrolase family-3 (GH3). We selected Cel3A because it has already been shown to enable *E. coli* to grow on cellobiose in minimal medium (12). CC_0968 was selected because *C. crescentus* is an α-proteobacterium like *Z. mobilis*. Past studies have successfully expressed *eglX* endoglucanase from *Pseudomonas fluorescens* var. *cellulosa* (later reclassified as *Cellvibrio japonicus sp. nov*) (13), *celZ* endoglucanase from *Erwinia chrysanthemi* (14), and endoglucanase from *Cellulomonas uda* CB4 (15) in *Z. mobilis.* Two cellulolytic enzymes from *Acidothermus cellulolyticus*, E1 and GH12, were expressed in *Z. mobilis* and their activities verified by a zymogram test with carboxymethyl cellulose (16). Similarly, β-glucosidase from *Ruminococcus albus* was tagged N-terminally with a signal peptide from the *Z. mobilis* periplasmic enzyme glucose-fructose oxidoreductase and overexpressed in *Z. mobilis*, which enabled fermentation of cellobiose to ethanol (9). However, these previous studies only demonstrated cellobiose conversion in resting cells, and heterologous gene expression in *Z. mobilis* was unable to produce significant growth in cellular biomass using cellobiose as a carbon source. Growth on oligosaccharides is crucial to enable genetic dissection of the *Z. mobilis* systems required for GH secretion and the use of selective pressure to improve GH production and section by *Z. mobilis*.

Here we report successful expression and secretion of a GH3 β-glucosidase encoded by *CC_0968* from *Caulobacter crescentus* at levels that enabled growth of *Z. mobilis* on cellobiose as a sole carbon source. Growth on cellobiose correlated with increased expression and secretion of GH3, which was induced by prolonged incubation (adaptation) in cellobiose or by exposure to sucrose medium. Proteomic analysis revealed that both cellobiose and sucrose adaptation included numerous changes to protein levels relative to growth on glucose that suggest a cellular remodeling program that enables GH secretion and possibly oligosaccharide transit across the outer membrane.

## RESULTS

### Heterologous expression of GHs in *Escherichia coli* and *Zymomonas mobilis*

Two GH-encoding genes, *cel3A* from *Cellvibrio japonicus* and *CC_0968* (encoding GH3) from *Caulobacter crescentus*, were cloned and expressed separately from an expression vector, pVector, which has an isopropyl β-D-1-thiogalactopyranoside (IPTG)-inducible T7A1 promoter driving GH expression, spectinomycin resistance gene, a Rhodobacter-derived broad host range origin of replication for *Z. mobilis*, and a pUC *ori* for *E. coli* (Fig 1A). *E. coli* DH5α and *Z. mobilis* ZM4 were transformed with pCel3A, pGH3, or control plasmid pVector and tested for growth in minimal medium containing either cellobiose or glucose as the sole carbon source. As expected, growth on cellobiose was not observed in either *E. coli* or *Z. mobilis* transformed with pVector (Figs 1B **and** E). However, for *E. coli* both pCel3A and pGH3 facilitated growth on cellobiose (Figs 1C **and** D). A corresponding effect was not observed for *Z. mobilis,* for which neither pCel3A nor pGH3 resulted in growth on cellobiose (Figs 1F **and** G). Both *E. coli* and *Z. mobilis* were able to grow on glucose when transformed with pVector, pCel3A, or pGH3, indicating that these plasmids do not negatively impact *Z. mobilis* viability (Figs 1B-G). Furthermore, *Z. mobilis* pVector and *Z. mobilis* pGH3 exhibited consistent growth in rich glucose medium (RMG) supplemented with spectinomycin and IPTG indicating that GH expression does not induce an inhibitory metabolic or protein synthesis burden in *Z. mobilis* pGH3 (**S1 Fig**).

**Figure 1.**
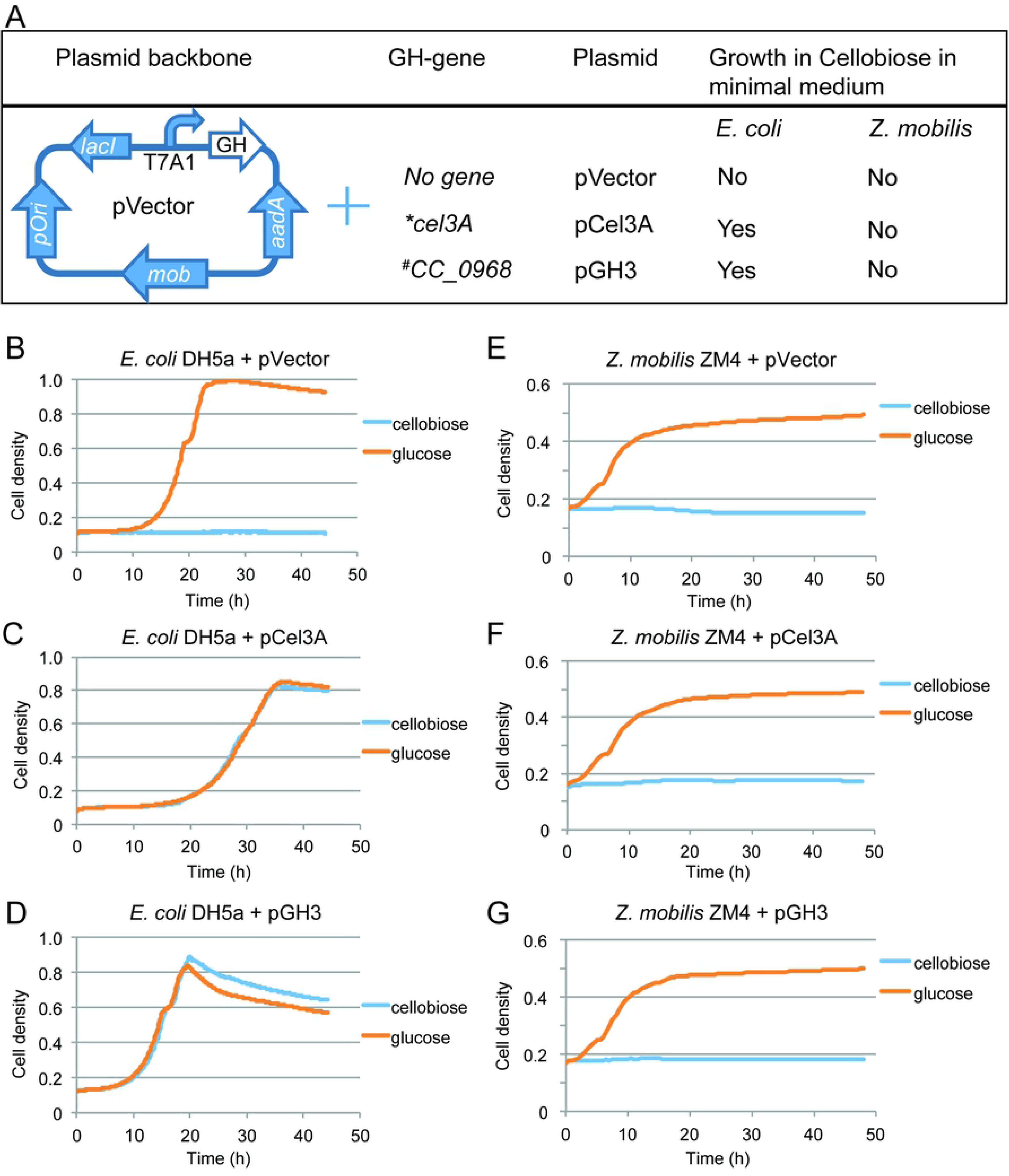
Expression of different glycosyl hydrolase (GH) genes from a common expression vector backbone and test of their effects on growth of *E. coli* DH5α in MOPS minimal medium (17) and *Z. mobilis* ZM4 in Zymomonas minimal medium (18) with glucose or cellobiose as a carbon source. (A) Expression of GH genes in a pVector backbone and a summary of the constructs and their effects on growth in minimal medium supplemented with cellobiose. (B-D) Growth of *E. coli* DH5α containing plasmids pVector or pCel3A or pGH3 in MOPS minimal medium supplied with 0.4% glucose or cellobiose. (E-G) Growth of *Z. mobilis* ZM4 containing plasmids pVector or pCel3A or pGH3 in a Zymomonas minimal medium containing 2% glucose or 2% cellobiose. The growth curves are averages of three replicates. *Gene from *Cellvibrio japonicus*, ^#^Gene from *Caulobacter crescentus*.

### *Non-native glycosyl hydrolases are readily secreted from* E. coli *but not* Z. mobilis

Bacteria express and localize proteins to various subcellular locations. Depending on the presence or absence of an N-terminal signal sequence and other features, proteins are either retained in the cytoplasm or localized to different cellular compartments. Secreted proteins may be targeted to inner membrane, outer membrane (OM), periplasm, or extracellular region depending on the accessibility of secretory apparatuses, the nature of signal sequences, or both (19). To determine why heterologous expression of GHs in *Z. mobilis* was unable to support growth on cellobiose, we tested for differences in expression and localization of the GHs between *E. coli* and *Z. mobilis*.

Extracellular and intracellular fractions of protein were separated as described previously (20). Briefly, equal numbers of cells (calculated based on culture volume and apparent OD) were centrifuged, and the cell pellets and supernatants used as the total cell and extracellular fractions. The periplasmic fraction was recovered after an osmotic shock of an equal portion of cells incubated in concentrated sucrose solution. The cytoplasmic fraction was obtained by lysis of osmotically shocked cells (see **Methods**). Using 4-methylumbelliferyl β-D-glucopyranoside (MUG), a fluorogenic analogue of cellobiose, GH activity was calculated in each protein fraction by measuring the accumulation of fluorescent reaction product over time using a 96-well plate reader (Table 1; **Materials and Methods**). Reported GH activity was normalized by the total protein concentration in the reaction.

**Table 1.** Localization and activity of glycosyl hydrolase expressed in *Z. mobilis* ZM4 and *E. coli* DH5α

| Fractions | *Z. mobilis* ZM4 in RMG medium |  |  | *E. coli* DH5α in LB medium |  |  |
| --- | --- | --- | --- | --- | --- | --- |
|  | ZM4+pVector | ZM4+pGH3 | % Activity | DH5α+pVector | DH5α+pGH3 | % Activity |
|  | GH activity(*U) | GH activity(*U) |  | GH activity(*U) | GH activity(*U) |  |
| Culture broth | nd | 2.0 ± 0.2 | 0.66 | nd | 15 ± 2 | 10.93 |
| Periplasm | nd | 210 ± 30 | 67.96 | nd | 68 ± 4 | 49.64 |
| Cytoplasm | nd | 80 ± 13 | 25.53 | nd | 12 ± 3 | 8.73 |
| Spheroplast | nd | 17 ± 1.2 | 5.48 | nd | 20 ± 2 | 14.41 |
| Whole pellets | nd | 1.2 ± 0.1 | 0.37 | nd | 22 ± 1 | 16.38 |

\*U, relative fluorescence signal produced per µg of protein per minute. nd, not detectable. RMG, rich medium glucose. LB, Luria-Bertani. Percentage activity was calculated from the ratio of activity of each fraction with respect to the total activity of all fractions.

We compared GH activity in *Z. mobilis* transformed with pGH3 encoding the *C. crescentus CC_0968* (called *Z. mobilis* GH3 hereafter), or with pVector, grown in rich medium glucose (RMG) to *E. coli* transformed with pGH3, or pVector, grown in Luria-Bertani (LB) medium. Among the isolated protein fractions of *Z. mobilis* GH3, GH activity was highest (∼68%) in the periplasmic fraction, with ∼99% of total GH activity localized to intracellular fractions (periplasm, spheroplast and cytoplasm). GH activity localization closely followed the localization predictions for *CC_0968* by LipoP 1.0 (21) and PSORTb (22) (**S3 Table**). Only a small fraction of activity (∼1% of total activity) was observed from the extracellular fraction (*i.e.*, culture medium; Table 1). *E. coli* pGH3 also exhibited about half of total GH activity in the periplasm but, in contrast to *Z. mobilis*, significant activity was also present in the culture medium and accessible in isolated whole cells (*i.e*., washed whole cells could hydrolyze assay substrate added to the resuspended cells; Table 1). These results suggest that extracellular or surface-accessible GH activity may be important for growth on cellobiose. Strains containing the pVector control plasmid did not exhibit GH activity in any fractions, indicating that *Z. mobilis* ZM4 and *E. coli* DH5α express little or no endogenous GH activity.

Although we detected expression and activity from both *C. crescentus* (pGH3) and *C. japonicus* (pCel3A) enzymes, subsequent experiments demonstrated that only pGH3 enabled adaptation of *Z. mobilis* for growth on cellobiose. Hence, we report here the subcellular GH activity for only GH3 from *C. crescentus*.

### Z. mobilis growth on cellobiose requires a long adaptation but not genetic mutation

After confirming GH activity in *Z. mobilis* GH3, we tested whether adaptation or selection for mutations could enable growth on cellobiose. *Z. mobilis* GH3 and a control strain were inoculated into rich medium 2% cellobiose (RMC) containing 0.4 mM IPTG at ∼0.05 apparent OD_600_. The cultures were incubated at 30°C and monitored for change in turbidity. No growth was evident for days one to three, whereas cells inoculated into rich medium 2% glucose (RMG) grew to saturation on day one. After three days, however, *Z. mobilis* GH3 started to grow in RMC (data not shown). To test whether the long lag could be eliminated by some cell growth to allow GH accumulation or other changes, we tested the same strains in RMC plus 0.05% glucose (RMCG). In the presence of 0.05% glucose, cells grew to an apparent OD_600_ of ∼0.13 at which point glucose was consumed from the medium. The cells then entered a long (∼48 hours) lag phase before again showing growth (Fig 2A). Cells grown in RMCG behaved similarly to cells grown in RMC such that a long lag phase was necessary before growth on cellobiose could occur.

**Figure 2.**
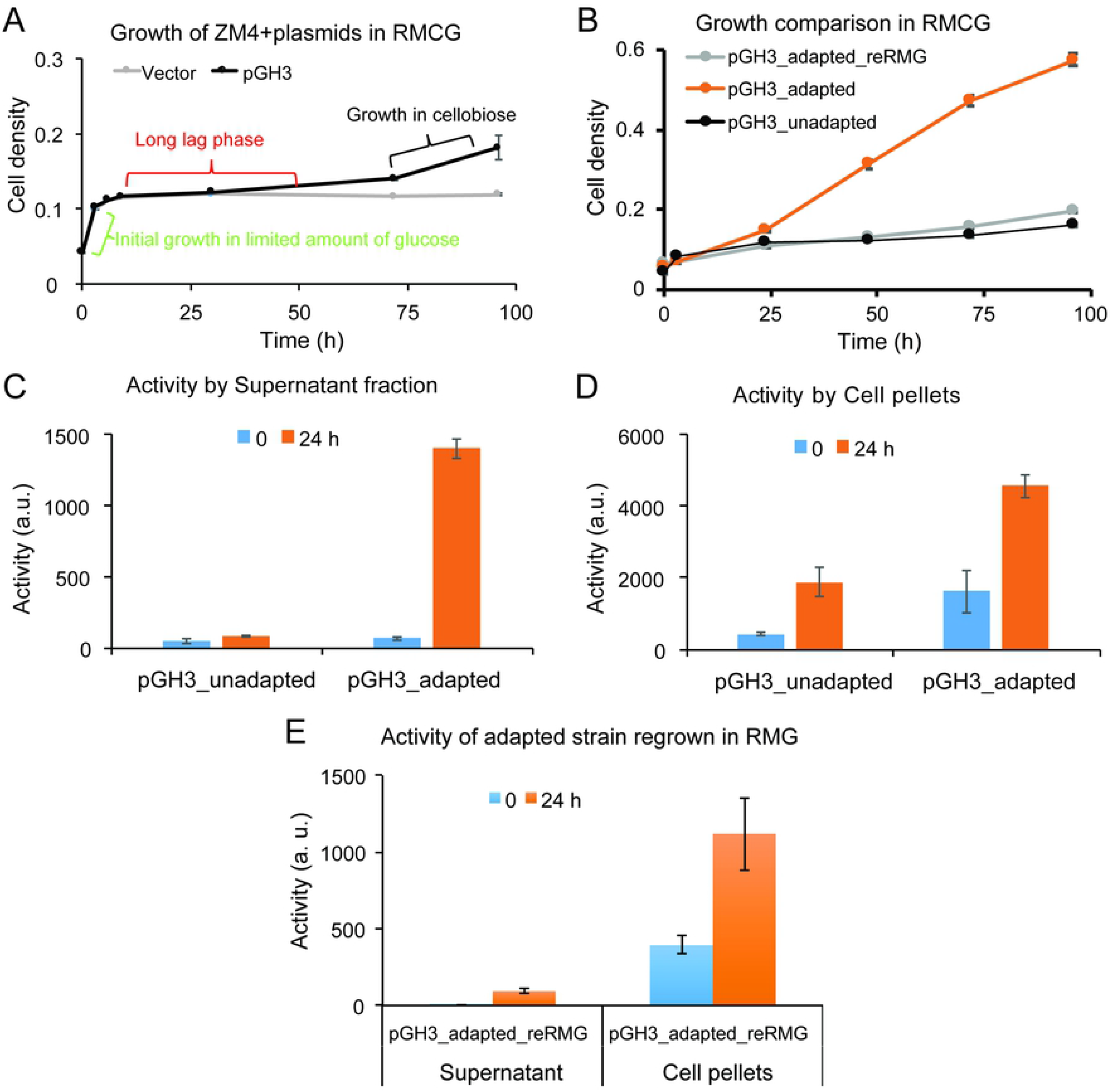
Growth and physiological changes induced by adaptation to cellobiose medium. (A) Growth of *Z. mobilis* ZM4 transformed with control pVector or pGH3 in 2% cellobiose medium with 0.05% glucose (RMCG). *Z. mobilis* GH3 growth in RMCG can be described in three stages: initial growth on glucose, a long lag phase, and growth on cellobiose. (B) Growth comparison of *Z. mobilis* GH3 in RMCG. Cells were either adapted to cellobiose, unadapted to cellobiose, or adapted to cellobiose and then regrown in RMG (reRMG). Extracellular (C) and whole-cell (D) GH activity for RMCG-adapted or -unadapted *Z. mobilis* GH3. (E) Extracellular and whole cell GH activity of RMCG-adapted *Z. mobilis* GH3 grown in RMG after adaptation before returning to RMCG. GH activity reported as relative fluorescence signal produced per min normalized by input cell number (apparent OD_600_). Error bars are standard deviations of biological triplicates.

To determine if growth on cellobiose was enabled by a permanent genetic alteration we grew RMC-adapted and -unadapted *Z. mobilis* GH3 in RMCG+0.4 mM IPTG. As a control, we also grew RMC-adapted *Z. mobilis* GH3 in RMG before returning cells to RMCG+0.4 mM IPTG (reRMG). Washed, RMC-adapted cells inoculated into RMCG no longer exhibited a long lag phase after consumption of glucose but instead continue growing on cellobiose (Fig 2B). RMC-unadapted cells exhibited a long lag phase after glucose consumption consistent with previous observations. Interestingly, the RMC-adapted cells grown in RMG before returning to RMCG lagged similarly to RMC-unadapted cells (Fig 2B) suggesting that growth on RMG reverted the cells back to a state that required a long lag phase before growth on cellobiose could resume. Based on these observations, we concluded that *Z. mobilis* GH3 adaptation to RMC cannot be attributed to a permanent genetic change. Instead we hypothesized that a slow remodeling of gene expression and cellular state occurs during adaptation (*i.e*., the long lag phase) that allows for growth on cellobiose.

We also tested the growth of *Z. mobilis* GH3 as colonies on solid agar medium containing RMC. Cells were struck out on RMC agar plates with or without 0.4 mM IPTG alongside control cells containing pVector and incubated for 3-6 days at 30 °C. After 3 days, adapted *Z. mobilis* pGH3 cells grew only in the presence of IPTG (**S2 Fig**), verifying that growth of *Z. mobilis* GH3 on cellobiose requires induction of GH3 synthesis. We also tested heterologous GH3 protein produced in *Z. mobilis* by comparing GH3 signal produced with and without IPTG. GH3 signal was increased with IPTG in all samples having GH3 protein in the plasmid (**S3 Fig**). Finally, to verify production of intact GH3, we visualized the induced GH3 protein by SDS-PAGE as a distinct, detectable band of ∼80 kDa (predicted MW of GH3 after removal of its signal sequence is 79652 Da) and estimated the extent of induction by densitometry (**S3 Fig**).

To understand the changes that allow *Z. mobilis* GH3 to grow on cellobiose, we used MUG to assay and compare extracellular (supernatant) and whole-cell (pellet) GH activity between adapted and unadapted cells. RMCG-adapted and unadapted cells were grown in RMC and samples were collected at zero and 24 hours. At zero hours both adapted and unadapted cells exhibited low extracellular GH activity. After 24 hours, adapted cells showed a marked increase in extracellular GH activity, whereas extracellular GH activity remained unchanged for unadapted cells (Fig 2C). Adapted whole cells exhibited higher GH activity than unadapted whole cells at both zero and 24 hours, with GH activity increasing several fold after 24 hours for both adapted and unadapted cells (Fig 2D). These results further indicate that GH secretion to the extracellular medium is important for growth on cellobiose. Adapted whole cells also displayed considerable levels of GH activity, suggesting that adaptation either increased permeability of the OM to the substrate (*i.e*., MUG or cellobiose) or increased display of GH3 on the cell surface. Taken together, either greater secretion of GH to the extracellular space, alteration of OMs, or both appear to contribute to growth on cellobiose.

We also measured extracellular and whole cell GH activity of the RMCG-adapted cells that were regrown in RMG before returning cells to RMC (reRMG). These reRMG cells showed low levels of extracellular GH activity and only moderate whole cell GH activity on par with unadapted cells (Fig 2E). Based on these findings, we concluded that adaption to RMC consists of a reversible remodeling of *Z. mobilis* rather than a genetic change that permanently altered the properties of the cells. When wild-type *Z. mobilis* ZM4 was transformed with plasmids recovered from RMC-adapted *Z. mobilis* GH3, the newly transformed strain behaved like unadapted cells, further indicating that the adaptation occurred in the host cell itself and not by mutation of the plasmid.

Having demonstrated that *Z. mobilis* GH3 can grow solely on cellobiose, we next measured cellobiose conversion to ethanol at successive times during adaptation and growth on cellobiose (0, 3, 6, 12, 24, 48, 96 and 168 hours after inoculation in RMCG medium). No cellobiose conversion occurred during the initial growth on glucose or during the approximately 48-hour adaptation period in *Z. mobilis* GH3. After 48 hours, *Z. mobilis* GH3 began consuming cellobiose as indicated by depletion of cellobiose from the medium (**S4A Fig**). Cellobiose depletion coincided with ethanol accumulation, reaching 3.4 ± 0.5 g/L after 168 hours of growth (**S4B Fig**). No cellobiose conversion was observed at any time for the control strain transformed with pVector (**S4A Fig**).

### Serial transfer of culture also enhanced cellobiose conversion

We also tested cellobiose adaptation of *Z. mobilis* GH3 by serial passage using *Z. mobilis* pVector as a control (Fig 3). The first passage was performed from RMG to RMC with 0.4 mM IPTG and all subsequent passages performed in RMC+IPTG (Fig 3A). To prevent carryover of extracellular GH between passages, cells were pelleted, washed, and resuspended in fresh medium. Growth of cells was monitored by measuring apparent OD_600_ and the supernatant collected at the beginning and end of each passage to measure cellobiose depletion and ethanol accumulation.

**Figure 3.**
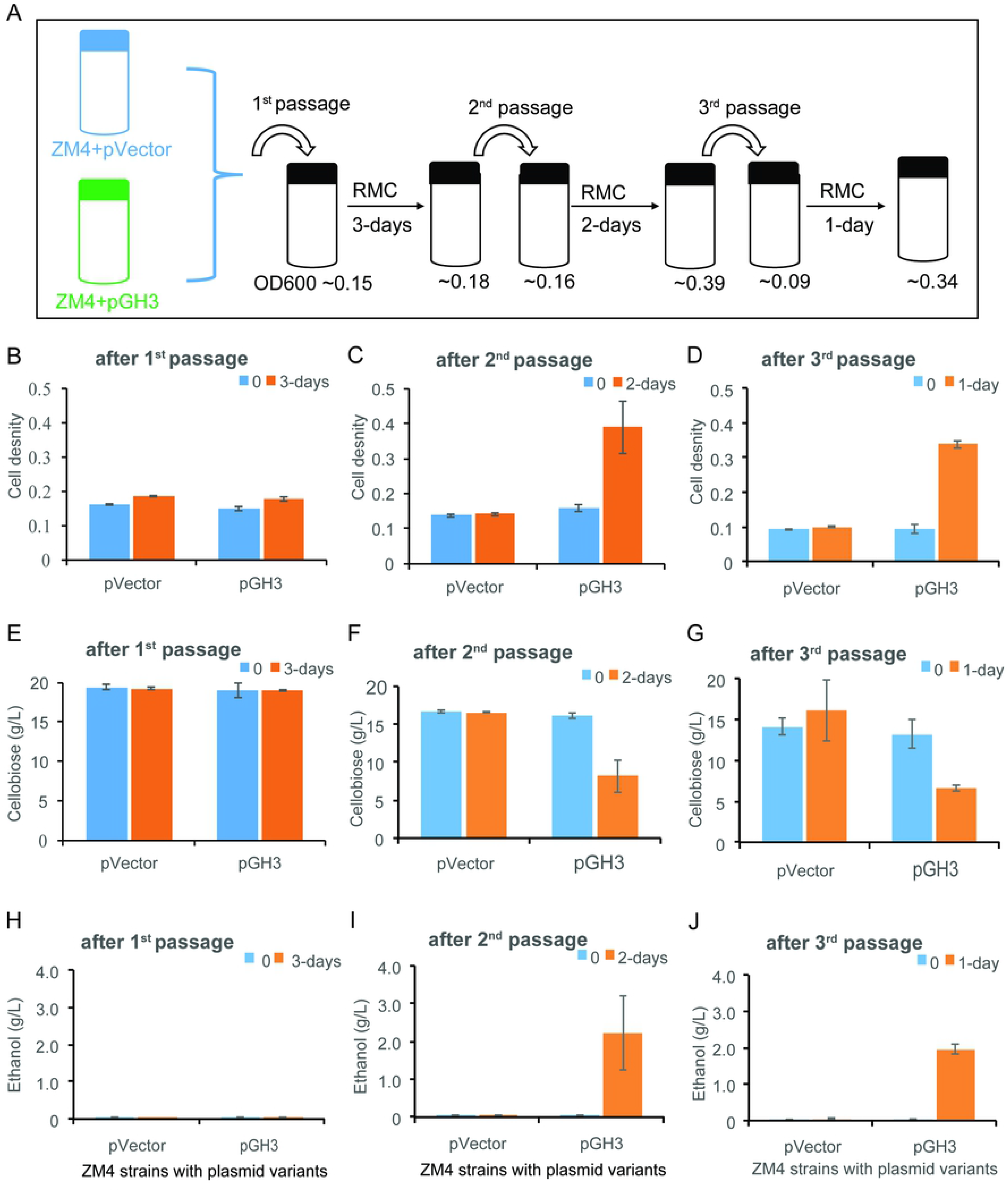
Growth adaptation by serial passage and its effect on growth in cellobiose and ethanol production. (A) Schematic representation of adaptation showing serial passages of *Z. mobilis* GH3 and pVector control. Apparent OD_600_ was measured for *Z. mobilis* GH3 at the end of each passage. (B, C, D). Growth of *Z. mobilis* GH3 and pVector control in RMC after first, second, and third passages. (E, F, G) Cellobiose conversion after first, second, and third passages, respectively. (H, I, J) Ethanol production after first, second, and third passages, respectively. Error bars are standard deviations of triplicate experiments.

After the first passage to RMC, little to no growth was observed over the course of 3 days for both *Z. mobilis* pVector and *Z. mobilis* GH3 (Fig 3B). After the second passage, *Z. mobilis* GH3 cell density more than doubled after two days whereas control cells did not grow (Fig 3C). *Z. mobilis* GH3 continued to grow robustly after the third passage with a doubling time of <24 hours (Fig 3D). During this serial passage growth of *Z. mobilis* GH3 on cellobiose was concomitant with cellobiose depletion from the medium (Figs 3E-G) and accumulation of ethanol (Figs 3H-J). From the serial passage experiment, we conclude that after the second passage *Z. mobilis* GH3 was able to utilize cellobiose for growth and ethanol production.

### Growth in sucrose medium enabled growth on cellobiose

Sucrose is a natural disaccharide substrate for *Z. mobilis*, catabolism of which depends on secreted sucrase(s) (23) and uncharacterized changes in cellular state. Thus, we hypothesized that exposure to sucrose might induce changes in *Z. mobilis* that enable growth on cellobiose. To test this hypothesis, we grew *Z. mobilis* GH3 in rich medium containing 2% sucrose (RMS) or RMG for 48 hours. The cells were then washed and inoculated into the fresh RMC containing 0.1% sucrose (RMCS) and growth was monitored (Fig 4A). We found that sucrose-grown cells resumed growth efficiently on cellobiose without a long lag phase, similar to RMC-adapted cells (Fig 2B), but neither RMG-grown *Z. mobilis* GH3 nor pVector control cells grew significantly. However, RMG-grown *Z. mobilis* GH3 did eventually resume growth after a lag, as seen previously (data not shown). This finding suggests that sucrose can induce changes in *Z. mobilis* that enable GH3-mediated growth on cellobiose. To investigate the effects of sucrose on GH activity, we assayed and compared GH activity of cellobiose-, sucrose-, and glucose-grown *Z. mobilis* GH3. We found that both cellobiose- and sucrose-grown cells exhibited higher extracellular and whole cell GH activity than glucose-grown cells (**S5 Fig**) consistent with our previous observations of RMC-adapted cells. These results suggest that growth on sucrose induces a cellular response in *Z. mobilis* that is similar to the remodeling that occurs during RMC adaptation.

**Figure 4.**
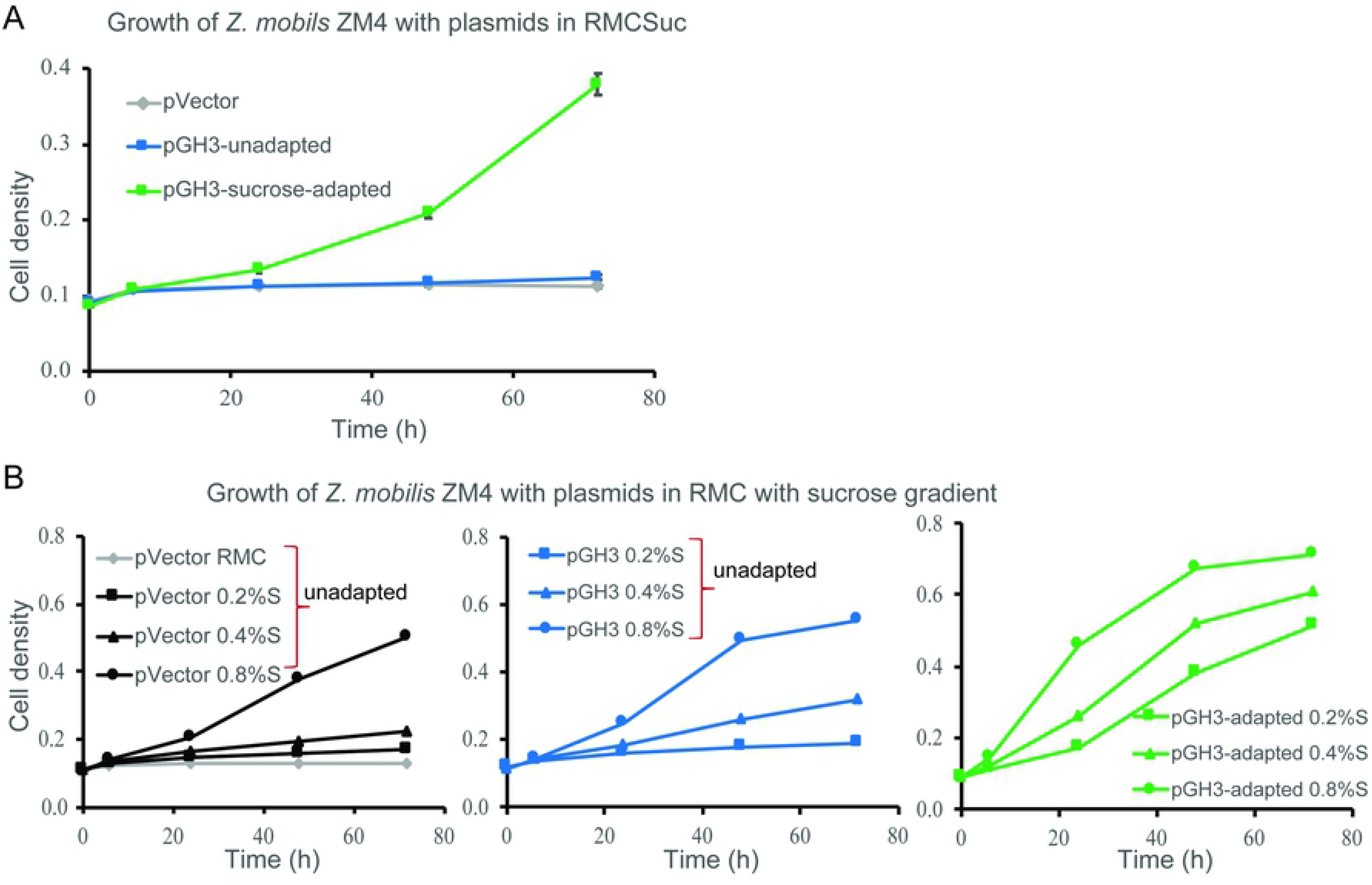
Sucrose adaptation allows *Z. mobilis* GH3 to grow on cellobiose. (A) Growth of sucrose-adapted *Z. mobilis* GH3, pVector control, and unadapted *Z. mobilis* GH3 in RMC supplemented with 0.1% sucrose (RMCSuc). (B) Growth of sucrose-adapted and unadapted *Z. mobilis* GH3 and pVector control, in RMC plus sucrose (0.2-0.8%). Gray line, *Z. mobilis* pVector RMC alone. Black lines, *Z. mobilis* + pVector in RMC+sucrose. Blue lines, *Z. mobilis* GH3 in RMC+sucrose. Green lines, sucrose-adapted *Z. mobilis* GH3 in RMC+sucrose.

To determine the minimum amount of sucrose needed to remodel cells for growth on disaccharides, we adapted *Z. mobilis* GH3 in RMC supplemented with 0.2 – 0.8% sucrose and compared growth in RMC+sucrose to unadapted cells and control pVector (Fig 4B). pVector control cells were only able to grow in RMC+0.8% sucrose, suggesting that 0.8% sucrose is the minimum amount of sucrose that will support growth of *Z. mobilis* in RMC medium (Fig 4B). After 72 hours, unadapted *Z. mobilis* GH3 grown in RMG before inoculating in RMC+sucrose showed little-to-no growth on RMC+0.2% sucrose and only modest growth on RMC+0.4% sucrose, the latter possibly supported by some cellobiose consumption (**S6 Fig**). Like pVector control cells, unadapted pGH3 cells grew in RMC+0.8% sucrose. However, we note that sucrose was significantly depleted from the medium by 48 hours compared to 72 hours for pVector control cells (**S6 Fig**). Continued growth of unadapted *Z. mobilis* GH3 in RMC 0.8% sucrose after 48 hours correlated with moderate consumption of cellobiose (**S6 Fig**). Sucrose-adapted *Z. mobilis* GH3, here defined as cells grown in RMS for 48 hours, were also inoculated into fresh RMC with increasing amounts of sucrose (*i.e*., RMC+0.2-0.8% sucrose). In each culture, the sucrose-adapted cells consumed almost all sucrose in the medium by 24 hours and after which cells continued to grow on cellobiose (Fig 4B **and S5C Fig**). These results suggest that 0.2% sucrose is sufficient to remodel *Z. mobilis* GH3 for growth on cellobiose, but that adaptation in higher sucrose concentrations will support more cell growth and greater rates of cellobiose consumption.

### Adaptation in cellobiose or sucrose medium remodeled Z. mobilis similarly

Adaptation to both cellobiose and sucrose similarly promote growth on cellobiose and induce increases in extracellular and whole cell GH activity. However, it is unclear what specific cellular changes occur in response to sucrose and cellobiose adaptation and what changes are needed for growth on cellobiose. To address this question, we compared protein levels of *Z. mobilis* GH3 adapted in cellobiose and sucrose and compared to unadapted cells grown in glucose (see **Methods**) using unlabeled mass spectrometry proteomics. Both extracellular and intracellular protein fractions were collected and analyzed. A total of 1539 proteins were identified from intracellular samples, representing >80% of annotated protein-coding genes in *Z. mobilis* ZM4 ATCC 31821 (5). A total of 1231 proteins were identified from extracellular samples, but most extracellular proteins (1215 out of 1231) overlapped with the intracellular fraction of proteins. This result suggests that many proteins detected in the growth medium (*i.e*., the extracellular fraction) are likely derived from the cytoplasm either by cell breakage or by incomplete separation of cells from the extracellular medium.

We observed greater similarity in intracellular protein levels (R^2^ = 0.37) between cellobiose- and sucrose-adapted cells when normalized to glucose-grown cells whereas extracellular protein levels were less similar across conditions (R^2^ = 0.17) (**S7 Fig**). Given our observations that extracellular GH activity increases during adaptation to cellobiose and that sucrase and levansucrase are known to be secreted from *Z. mobilis* (23) in response to sucrose, we looked at levels of secretion-related proteins in both sucrose- and cellobiose-adapted cells. Notably, the levels of a majority of annotated transport and secretion-related proteins increased in both sucrose- and cellobiose-adapted cells (Fig 5, **S7 Fig**). Interestingly, GH3 CC_0968 was also upregulated in both extracellular and intracellular fractions in both cellobiose and sucrose media (Fig 5, **S8 Fig**) despite a consistent amount of IPTG in the glucose, sucrose, and cellobiose media. This increase in GH3 expression may be explained by an increase in pGH3 plasmid copy number, reduced turnover of GH3 after secretion to the extracellular medium, or both.

**Figure 5.**
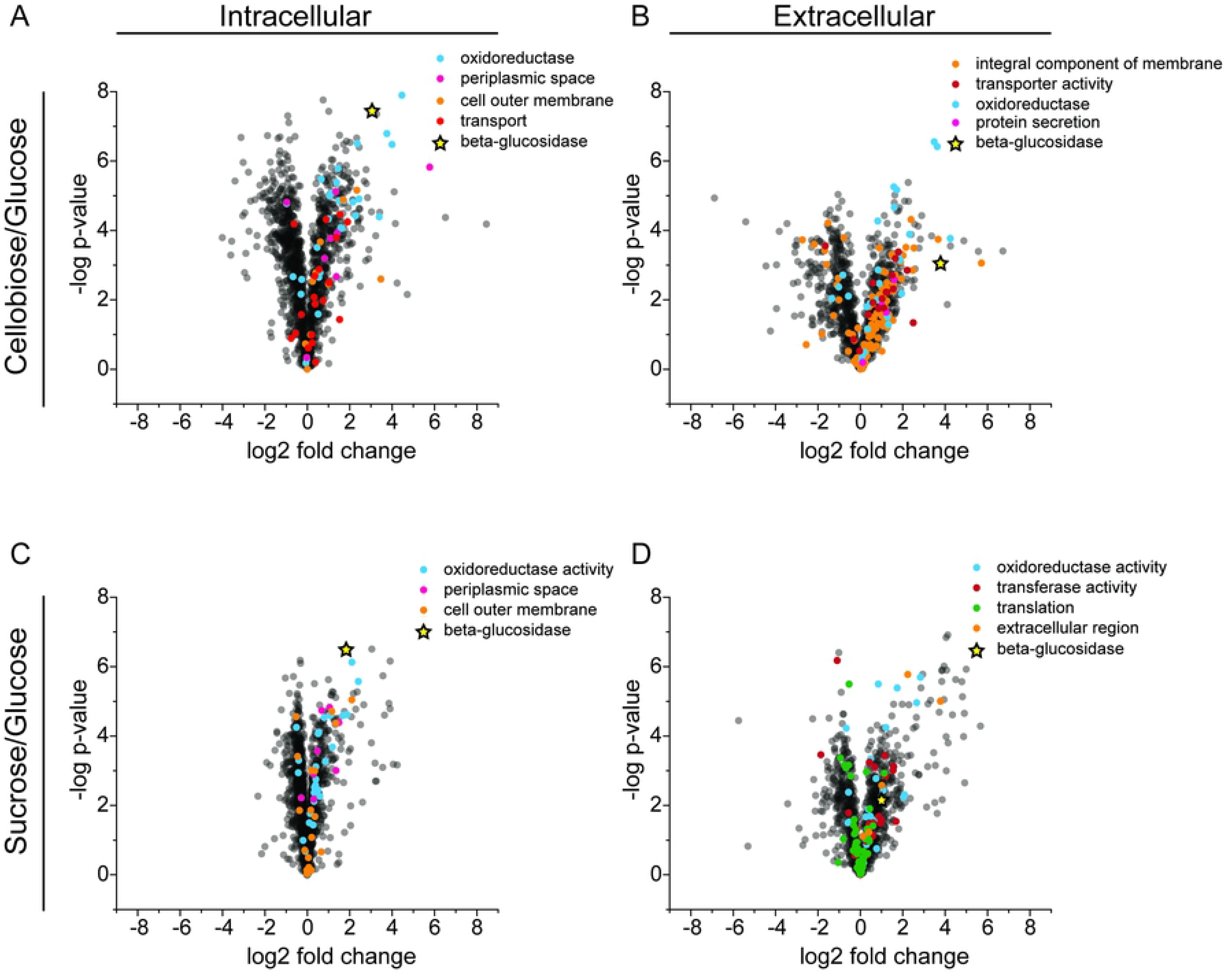
Volcano plots of Gene Ontology (GO) enrichment analysis showing differential expression of GO term proteins. The upper panels show (A) intracellular and (B) extracellular proteomics for cellobiose-adapted *Z. mobilis* GH3 compared to unadapted strain. The lower panels show (C) intracellular and (D) extracellular proteomics for sucrose-adapted *Z. mobilis* GH3 compared to unadapted strain. All enriched GO term proteins are indicated with spheres of distinct colors and β-glucosidase upregulation is shown in highlighted star.

To understand the nature of protein remodeling during sucrose and cellobiose adaptation more completely, we performed K-means clustering on normalized log2-fold change values (normalized to glucose-grown cells) for proteins that were measured in both the intracellular and extracellular fractions (1199 proteins in total). In total, 50 clusters were produced revealing similar remodeling patterns between sucrose- and cellobiose-adapted cells (Fig 6). Two prominent clusters are present in which protein levels are similarly downregulated or upregulated in sucrose- and cellobiose-adapted cells. We performed Gene Ontology (GO) enrichment analysis of KEGG pathways (24) for each cluster and identified several pathways that were statically enriched across six clusters. Of the clusters primarily comprised of upregulated proteins, several transport-related and cell membrane pathways were enriched such as the ABC transporter, integral membrane component, receptor activity, transported activity, and cell outer membrane pathways (Figs 5 and **6**). These results are consistent with GO-term enrichment analysis of differential protein levels in each sample compared to glucose-grown cells where GO terms related to stress (oxidoreductase), secretion, and transport (periplasmic space, OM, transport, and protein secretion) were enriched in both intra- and extracellular fractions of cellobiose- and sucrose-adapted strains (**S4 Table**). Within the large cluster of primarily downregulated proteins several growth-related pathways were enriched including the ATP synthase, ribosome, translation, and purine and pyrimidine biosynthesis pathways (Fig 6). A decrease in growth-related proteins can be attributed to the reduced growth rate of cells metabolizing cellobiose or sucrose relative to cells growing on glucose.

**Figure 6.**
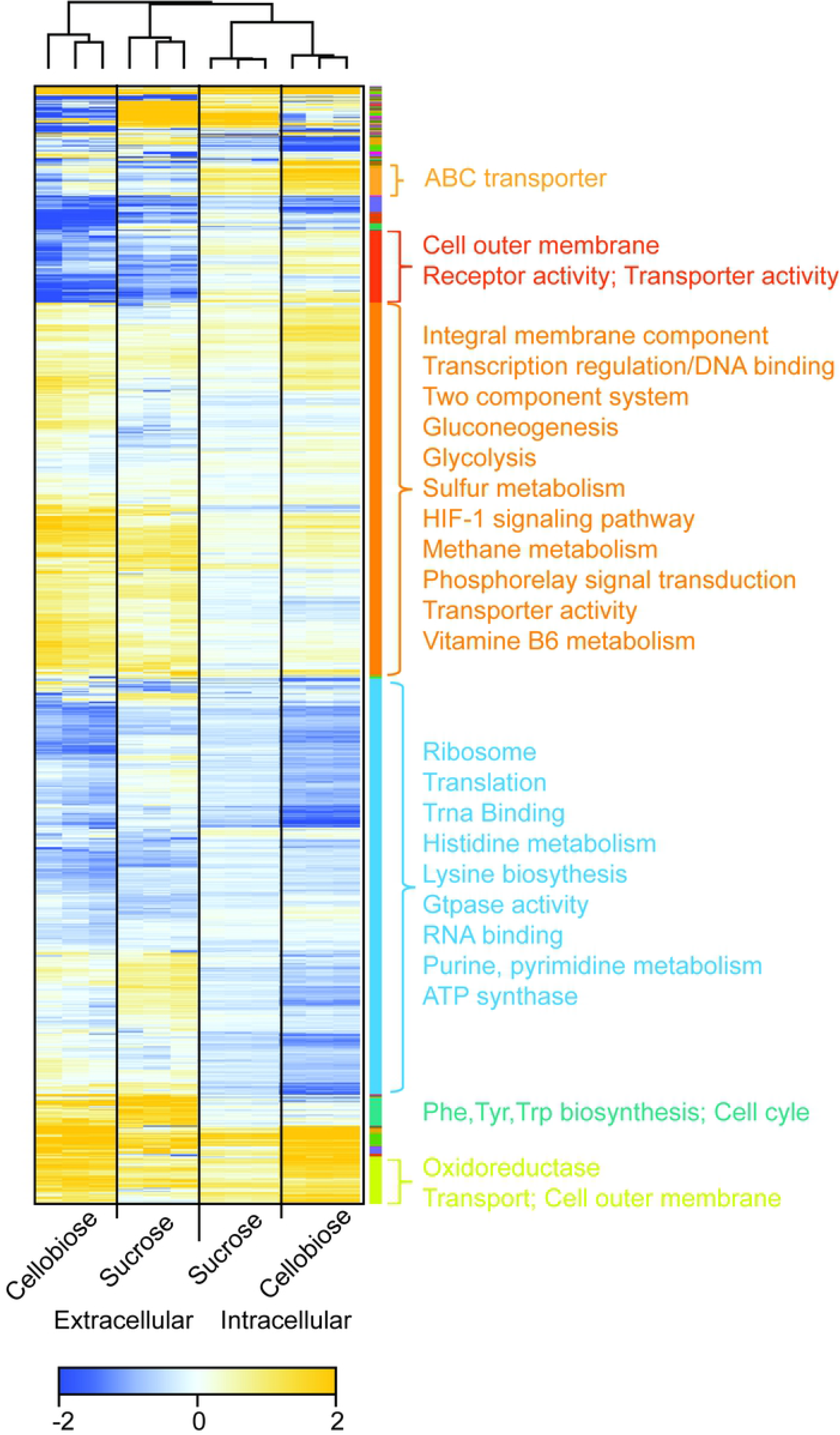
Heat map displaying hierarchical clustering of control-normalized log2 fold changes of 1199 molecules quantified across 12 replicates. Both row-wise and column-wise clustering was performed using Euclidean distance and average linkage calculations. 50 distinct hierarchical clusters are represented in the color bar shown alongside the heat map and only the significant protein families are indicated.

Despite the similarities in protein expression between cellobiose- and sucrose-adapted cells, we observed sets of proteins uniquely differentially expressed between the two conditions (**S7 Fig**). Of note, we observed a disparity in the expression patterns of SacB (extracellular levansucrase) and SacC (extracellular sucrase) (25–27). Although SacB was upregulated in both cellobiose and sucrose adapted cells (**S1 Data**), SacC expression was only upregulated in sucrose-adapted cells. Previous work has shown that SacB and SacC can be expressed as both a bicistronic transcript and individually as monocistronic transcripts with SacC under control of a strong promoter (28). Our results indicate that SacC upregulation is dependent on a sucrose-specific signal whereas SacB monocistronic expression can be activated in a sucrose-independent manner.

## DISCUSSION

Our study of xenogeneic GH-enabled growth of *Z. mobilis* on cellobiose and its conversion to ethanol revealed two key discoveries. First, even though high expression of a β-glucosidase from a closely related α-proteobacterium *C. crescentus* occurs relatively quickly (within 12 hours) in rich medium glucose, *Z. mobilis* GH3 depends on a transient adaptation period of several days before growth on cellobiose and high-flux ethanol production becomes possible. Second, growing *Z. mobilis* on sucrose induces cellular changes similar to cellobiose adaptation that also allow *Z. mobilis* GH3 to readily grow on cellobiose without a long lag phase.

### Cellobiose adaptation coincides with extensive changes to the *Z. mobilis* envelope

In this study, we showed that adaptation of *Z. mobilis* GH3 to cellobiose medium correlated with extensive changes to its cell envelope which coincided with an increased presence of GH3 in the extracellular medium. When grown in RMG, GH3 was located mostly in the periplasm and cytoplasm with only a small fraction of GH3 secreted to the culture medium (**Table 1**). These initial low levels of extracellular or periplasmic GH were insufficient to generate enough glucose for growth and the strain required an approximately 72-hour lag phase before cells became adapted and slow growth could commence (Fig 2A). Growth on cellobiose consistently coincided with increased GH activity in the extracellular or outer membrane space (**Table 1,** Fig 2**, S5 Fig**) underscoring the importance of GH localization for cellobiose conversion. Once cells expressing GH3 were adapted to cellobiose, they could continue to grow without a long lag when placed in fresh cellobiose medium. However, this adaptation to cellobiose was reversible, and when cellobiose-adapted cells were regrown in RMG they would again require a long lag phase (Fig 2B) in cellobiose medium before growth could occur. Thus, cellobiose adaptation consisted of a reversible change to the cellular composition that was not conferred by a stable genetic adaptation.

Our proteomics results revealed that adaptation to cellobiose changed the protein composition of *Z. mobilis* GH3 compared to a glucose-grown control. The protein expression patterns clustered into groups of proteins of related functions in both the intracellular and extracellular fractions (Fig 6). Notably, proteins related to secretion and transport were upregulated (Fig 5) which we hypothesize lead to increased secretion of GH3, increased display of GH3 on the outer membrane or entry of substrates (cellobiose) into the periplasm, or all of the above (Fig 2D).

### Cellobiose and sucrose adaptation may be comprised of a native scavenging response to glucose depletion

Similar cellular changes were observed during adaptation on cellobiose and adaptation to sucrose, a native disaccharide substrate in *Z. mobilis*. This similarity is apparent in the protein level changes in cellobiose- and sucrose-adapted cells, including increased extracellular GH3, and by the observation that adaptation to sucrose enables growth on cellobiose. Although sucrose is a natural substrate for *Z. mobilis*, we also observed a long lag phase before cells began growing on sucrose. Given these findings, we hypothesize that the changes in cellular envelope and protein secretion that accompany both sucrose and cellobiose adaptation are part of a generalized scavenging response to glucose-depleted conditions. The changes in protein secretion and envelope composition during this scavenging response could improve acquisition of nutrients by *Z. mobilis* from the extracellular medium. Should a suitable substrate be found while scavenging, then substrate-specific and possibly more energy intensive responses would be activated as we observed with sucrose-specific upregulation of SacC. In the case of cellobiose adaptation this cellular remodeling increased GH3 secretion or surface display of cells incubated with cellobiose and ultimately enabled cells to grow on cellobiose.

### Prospects for engineering Z. mobilis for broader utilization of oligosaccharides

*Z. mobilis* is being developed as a platform for the conversion of lignocellulosic hydrolysates to biofuels and bioproducts. Complete utilization of oligosaccharides will improve product yields and economic feasibility of lignocellulosic biomass conversion. Our findings highlight important challenges to engineering broader oligosaccharide utilization in *Z. mobilis*. First and foremost, we lack a comprehensive understanding of native secretion and transport pathways in *Z. mobilis*. Broadening the substrate specificity to oligosaccharides beyond cellobiose will require a more complete understanding of the rules governing both GH secretion and oligosaccharide entry to the periplasm in *Z. mobilis.* It is clear from our work and the work of others (29, 30) that GH localization and substrate accessibility is crucially important. Likewise, the impact of species origin on heterologous gene expression in *Z. mobilis* may not be fully appreciated. That expression of *celA* from *C. japonicus* did not enable *Z. mobilis* to grow on cellobiose, even when allowing time for adaptation, suggests that important differences exist in how xenogeneic glycosyl hydrolases are recognized by and interact with endogenous *Z. mobilis* pathways.

Although GH3 expression allows *Z. mobilis* to metabolize cellobiose, the long adaptation time required to remodel *Z. mobilis* for growth on cellobiose is not practical for industrial applications. Continuous fermentations may be a suitable option given that cellobiose adaptation is retained upon transfer to fresh RMC. However, this option does not eliminate the initial long adaptation time. Further mechanistic dissection of cellobiose and sucrose adaptation is needed to identify key regulators governing adaptation (e.g., the proposed scavenging response). With greater understanding of the basic mechanisms, it is plausible that GH-expressing strains could be engineered by rewiring the natural response to sucrose to induce the necessary cellular changes to support oligosaccharide metabolism and eliminate the need for a long adaptation to achieve cellular remodeling.

## MATERIALS AND METHODS

### Strains, plasmids and culture conditions

*Zymomonas mobilis* ZM4 (ATCC #31821), *Escherichia coli* DH10B (Invitrogen, Carlsbad, CA, USA) and *E. coli* DH5α (New England BioLabs, Ipswich, MA, USA) were used in this study. The *E. coli* DH10B strain was used for cloning and *E. coli* DH5α was used for expressing the recombinant plasmids. Unless otherwise specified, all the *E. coli* strains were grown in LB or MOPS minimal medium (17) at 37°C with shaking. *Z. mobilis* strains were grown in rich medium containing 1% yeast extract, 15 mM KH_2_PO_4_ plus 2% glucose (RMG), 2% cellobiose (RMC), varying concentrations of sucrose (RMS), or in *Zymomonas* minimal medium (ZMM) (18) at 30°C without shaking. Plasmid pIND4-spec, a derivative of the *Rhodobacter*-derived, broad-host-range shuttle vector pIND4 (31) in which the kanamycin-resistance gene was replaced with a spectinomycin resistance gene, was used to clone and express glycoside hydrolases. Spectinomycin concentration at 50 µg ml^-1^ for *E. coli* and 100 µg ml^-1^ for *Z. mobilis* were used.

### Plasmid construction

All oligonucleotide primers used for cloning are listed in **S1 Table** and were obtained from Integrated DNA Technologies (Coralville, IA, USA). For plasmid construction, the vector backbone and gene fragments were amplified by PCR using Q5 DNA polymerase (New England Biolabs, Ipswich, MA, USA) following the manufacturer’s protocol. DNA fragments were purified by agarose gel electrophoresis and assembled using Gibson Assembly mix (New England Biolabs, Ipswich, MA, USA) following the manufacturer’s protocol. All plasmids used in this study were verified by DNA sequencing and restriction enzyme digestion analysis and are listed in **S2 Table**.

The plasmids pCel3A and pGH3 were constructed by cloning the respective glycosyl hydrolases encoded by *cel3A* from *Cellvibrio japonicus* and *CC_0968* from *Caulobacter crescentus* in pIND4-spec (pVector). The predicted localizations of Cel3A and CC_0968 are shown in **S3 Table**.

### Transformations of *E. coli* and *Z. mobilis* cells

Electro-competent cells were prepared following standard protocol (32). About 100 ng of plasmid DNA or Gibson assembly mixture was used to transform ∼10^9^ cells in 50 μl of *E. coli* competent cells in 10% glycerol. Electroporation of *E. coli* was performed with a Bio-Rad gene pulser with a setting of 200Ω, 25 μF, and 1.75 kV in a 0.1 cm cuvette. Immediately after electroporation, 1 ml SOC medium was added and cells were incubated at 37°C for 1 hour; 100 μl of the recovered cells were then spread on LB agar containing appropriate antibiotic for selection and the plates were incubated at 37°C overnight.

For *Z. mobilis* transformation, ∼1 μg plasmid DNA was used to transform ∼10^9^ cells in 50 μl in 10% glycerol. Type 1 restriction inhibitor (1 µl; Epicentre) was added to the plasmid DNA prior to mixing with competent cells. Electroporation of *Z. mobilis* was performed with a Bio-Rad gene pulser with a setting of 200Ω, 25 μF, and 1.6 kV in a 0.1 cm cuvette. Immediately after electroporation, 1 ml recovery broth (5 g glucose/L, 10 g yeast extract/L, 5 g tryptone/L, 2.5 g (NH_4_)_2_SO_4_/L, 0.2 g KH_2_PO_4_/L, and 0.25 g MgSO_4_•7H_2_O/L) was added and the cells were incubated for 2-3 hours at 30°C. The recovered cells were spread on RMG-agar containing the appropriate antibiotic for selection and the plates were incubated at 30°C for 2-4 days to obtain transformed colonies.

### Extraction of cellular and subcellular fractions, protein quantification, and activity assay

For activity measurements, seed cultures were prepared by overnight cultivation in the desired conditions. Five ml of LB or RMG supplemented with required concentration of spectinomycin were inoculated with seed cultures of *E. coli* and *Z. mobilis*, respectively, and incubated until the apparent OD_600_ reached ∼0.4. For protein induction, 0.2 mM IPTG was added to both cultures and incubation was continued overnight. Extracellular and intracellular fractions (supernatant, periplasm, cytoplasm, spheroplast and whole cells) were prepared from the same number of cells quantified by measuring apparent OD_600_ and adjusting the volume to obtain an apparent OD_600_ of 1.5.

The supernatant fraction (supernatant/culture medium) was obtained by centrifugation of the cell culture at 20,000 x *g* for 3 minutes at 4°C. Spheroplasts were prepared by the osmotic shock protocol (20). Briefly, after removing the culture medium as supernatant, the pellets were resuspended in 500 μl of 20 mM Tris-Cl pH 8.0, 2.5 mM EDTA, 20% (w/v) sucrose and incubated on ice for 10 minutes. The sample was then centrifuged for 3 minutes at 20,000 x *g* at 4°C and supernatant was discarded. The pellets were resuspended in 300 μl of sterile ice-cold water and incubated in ice for 10 minutes. After centrifugation at 20,000 x *g* for 3 minutes at 4°C, the supernatant was collected as periplasmic fraction (periplasm) and the remaining pellets (spheroplasts) were resuspended in 300 μl sterile ice-cold water. For preparation of cytoplasmic fraction, spheroplasts were treated with 50 μl Popculture (Novagen, Madison, WI, USA), 50 μl lysozyme solution (10 mg/ml) and 200 μl sterile water and incubated at 30°C for 30 minutes. After centrifugation, the supernatant was collected as cytoplasmic fraction (cytoplasm). Whole cells were prepared by removing culture medium by centrifugation and resuspension in 300 μl sterile water.

Protein concentration was measured by using Bicinchoninic Acid (BCA) assay (Thermo Scientific) following the manufacturer’s protocol. Twenty microliter cell samples and standards of diluted bovine serum albumin (BSA) were transferred to clean 1.5 ml microtubes. A no protein control was also included. Cold (−20°C) acetone (80 µl) was added and the sample was vortexed vigorously and incubated at −20°C for 1 hour. The proteins were pelleted by centrifugation at 4°C at 15,000 x *g* for 15 minutes. The supernatant was then carefully removed and discarded. Protein pellets were washed with 100 µl of cold 100% acetone by adding it around the walls and spinning at 4°C 15,000 x *g* for 2 minutes. After removal of supernatant, the pellets were dried *in vacuo* at room temperature for 10 minutes. The dried pellets were resuspended in 20 µl of 2% SDS, 9 mM Tris-Cl pH 8.0 and incubated at 70°C for 10 minutes. Protein concentration was measured following the manufacturer’s protocol. Samples were incubated at 37°C for 30 minutes to 2 hours. Absorbance was measured at 562 nm in a Tecan M1000 plate reader (Tecan Group Ltd., Männedorf, Switzerland) and protein concentration was determined by comparison to BSA standards.

The glycosyl hydrolase activity assay was performed by adding 20 or 25 μl of protein sample to 50 μl of 2 mM 4-methylumbelliferyl β-D-glucopyranoside (MUG; Sigma-Aldrich, St. Louis, MO, USA) in a 96-well plate. The reaction was monitored in a Tecan M1000 plate reader with fluorescence excitation at 365 nm and emission at 455 nm for 180 minutes with readings every 5 minutes. The fluorescence produced was plotted as a function of time and the enzyme activity was determined from the slope of this plot. The activity value was normalized by the amount of protein present in the reaction.

### Growth adaptation

Triplicate samples of *Z. mobilis* GH3 (expressing *CC_0968*) and the pVector control were grown overnight in 5 ml RMG medium containing spectinomycin at 30°C. One ml of each sample at apparent OD_600_ ∼1.0 was centrifuged and washed twice with sterile RMC medium. Washed cell pellets were then resuspended in a culture tube with 10 ml RMC medium containing spectinomycin and 0.4 mM IPTG. Culture tubes were then incubated at 30°C without shaking. Growth was monitored initially after 3, 6, 12 and 24 hours, and then every 24 hours. After a significant growth was seen for *Z. mobilis* GH3, cells were collected, washed with RMC medium, and resuspended again in RMC with spectinomycin and IPTG. This process was repeated with RMC medium supplemented with 0.05% glucose (RMCG) and the data are presented in this study (Fig 2A). To evaluate whether the adaptation was due to permanent genetic change, cellobiose-adapted *Z. mobilis* GH3 was regrown in RMG medium (reRMG) and transferred back to fresh RMCG medium after washing. Growth of three types of *Z. mobilis* GH3 strains, unadapted, adapted and adapted regrown in RMG (reRMG), were then compared in identical conditions (Fig 2B).

A serial passage experiment also was performed using *Z. mobilis* GH3 and pVector (Fig 3). The cells were grown in RMG overnight, collected by centrifugation and washed with RMC to remove residual glucose. The cells were then resuspended in RMC containing spectinomycin and 0.4 mM IPTG to an apparent OD_600_ of ∼0.1 and incubated at 30°C for 3 days (first passage). After 3 days, apparent OD_600_ was measured and the second passage was performed after centrifugation and resuspension of the cell pellets in fresh RMC-spectinomycin-PTG medium at a similar starting apparent OD_600_. Incubation was continued for 2 days and then the third passage was performed after similar recovery by centrifugation and resuspension of the cell pellets in fresh RMC-spectinomycin-IPTG medium. For the third passage, incubation was continued for one day. Cultures were then quantified for apparent OD_600_ and stored for metabolite analysis by HPLC.

For adaptation in sucrose medium, *Z. mobilis* GH3 was grown in rich medium containing 2% sucrose (RMS) at 30°C for 48 hours without shaking. After 48h, cells were collected by centrifugation at 5000 x *g* for 5 minutes at room temperature. Supernatant was discarded and pellets were washed with sterile deionized water. Finally, the pellets were resuspended in a medium where the subsequent culture was to be done.

### Growth and activity measurement from adapted *vs* unadapted culture

Adapted cultures were derived from cells grown in RMCG for 24 hours. Unadapted cultures were derived from cells grown in RMG for 12 hours. After centrifugation, the pellets were washed with RMC and resuspended in RMCG medium containing spectinomycin and IPTG. For growth measurement, samples were removed at time intervals, apparent OD_600_ was recorded and cells were stored at −20°C prior to metabolite analysis. For GH activity measurements, the required volume of cells was withdrawn from the culture tube, centrifuged at 20,000 x *g* for 5 minutes. The supernatant was transferred to a fresh tube to assay for extracellular GH activity and the pellets were washed with water and resuspended in water at a calculated volume to give an equivalent number of cells in all samples. For GH activity measurement, 20 or 25 µl of extracellular or pellet fractions were transferred to a 96-well plate in triplicate, 50 µl of 2 mM MUG was added, and the readings in a Tecan 1000 plate reader were immediately started (fluorescence excitation at 365 nm and emission of 455 nm and, for cell samples, cell density at 600 nm). Normalization was performed by dividing the fluorescence value with corresponding apparent OD_600_.

### SDS PAGE and GH3 protein signal measurement

For analysis of GH3 protein induction, both adapted and unadapted *Z. mobilis* strains with plasmids pGH3 or pGH3T were grown in replicate in RMG supplemented with appropriate concentration of antibiotics (**S3 Fig**). The plasmid pGH3T encodes, in addition to GH3, a *C. crescentus* TonB receptor for cellobiose (*CC_0970*) that was found to have little or no effect on cellobiose utilization in *Z. mobilis.* To one replicate, IPTG was added to a final concentration of mM when the culture reached apparent OD_600_ of ∼0.4 and growth was continued for 24 hours. Final apparent OD_600_ was measured for all samples. Approximately equal numbers of cells equivalent to one ml of apparent OD_600_ ∼3.0 were centrifuged at 10,000 x *g* for 5 minutes. The pellets were resuspended in 20 µl of 1x SDS loading solution (62 mM Tris·Cl, pH6.8, 2% w/v SDS, 10% v/v glycerol, 5% v/v β-mercaptoethanol, 0.05% w/v bromphenol blue), incubated at 98°C for 10 minutes, and then immediately cooled on ice for 5 minutes. The samples were centrifuged at 10,000 x *g* for 2 minutes and 10 µl portions of the supernatants were loaded on a 4-12% Tris-Glycine slab gel connected to Tris-Glycine SDS running buffer (Invitrogen). The gel was electrophoresed at 200 volts for 1 hour, stained with Coomassie Brilliant Blue R-250 solution (BioRad, #1610436), imaged using a white light transilluminator and a CCD camera equipped with an 595 ± 55 nm bandpass filter (Fluorichem 8000; Protein Simple, Inc.), and then quantified using Imagequant software (GE Healthcare).

### Quantification of cellobiose conversion and ethanol production

Samples were withdrawn after apparent OD_600_ measurement and stored at −20°C until all required time points were collected. The frozen samples were thawed at room temperature and vortexed and centrifuged prior to subsampling. 100 µl of the samples were transferred to labeled 1.5 ml autosampler vials and 900 µl of pure water was added to each vial and mixed properly. The vials were capped and placed in a 4°C cooled autosampler tray. Fifty µL were injected to an Agilent 1260 Infinity HPLC system with a quaternary pump, vacuum degasser, and refractive index detector (Agilent Technologies, Inc., Palo Alto, CA) and separated on an Aminex HPX-87H with Cation-H guard column (BioRad, Inc., Hercules, CA, USA) 300×7.8mm, cat #125-0140. The mobile phase was 0.02 N H_2_SO_4_ was used at a flow rate of 0.5 ml/min, and both column and detector temperatures were maintained at 50°C. Data were analyzed using ChemStation C.01.06 software (Agilent Technologies, Inc., Palo Alto, CA, USA). The metabolites of interest (cellobiose, glucose, and ethanol) were analyzed and quantified using standard calibration curve prepared from the respective pure compounds obtained from Sigma-Aldrich (St. Louis, MO, USA).

### Proteomics analysis of cellobiose- and sucrose-adapted vs unadapted *Z. mobilis* GH3

*Z. mobilis* GH3 was grown in RMG, RMC, or rich medium + 2% sucrose (RMS) until the apparent OD_600_ reached to late log phase. Cells were harvested and extracellular and intracellular fractions were collected by centrifugation at 20,000 x *g* for 5 minutes at 4°C. Proteins were digested, analyzed by LC-MS/MS, and peptide identity was verified with *Z. mobilis* genome peptide library as described below.

#### Lysis and digestion

Cells were lysed by suspension in 6 M guanidine hydrochloride (GnHCl), followed by addition of methanol to 90%. Samples were centrifuged at 15,000 x *g* for 5 minutes at 4°C. Supernatants were discarded and pellets were allowed to air dry for ∼5 minutes. Pellets were resuspended in 200 µL 8 M urea, 100 mM Tris pH 8.0, 10 mM (tris(2-carboxyethyl)phosphine) (TCEP), and 40 mM chloroacetamide, then diluted to 2 M urea in 50 mM Tris pH 8. Trypsin was added at an estimated 50:1 ratio, and samples were incubated overnight at ambient temperature. Each sample was desalted over a PS-DVB solid phase extraction cartridge and dried *in vacuo*. Peptide mass was assayed with a peptide colorimetric assay.

#### Liquid chromatography with tandem mass spectrometry (LC-MS/MS)

For each analysis, 2 µg of peptides were loaded onto a 75 µm inner diameter, 30 cm long capillary with an imbedded electrospray emitter and packed with 1.7 µm C18 BEH stationary phase. The mobile phases used were A: 0.2% formic acid and B: 0.2% formic acid in 70% acetonitrile. Peptides were eluted with an increasing gradient of acetonitrile from 0% to 53% B over 75 minutes followed by a 5 minute 100% B wash and a 10 minute equilibration in 0% B.

Eluting peptides were analyzed with an Orbitrap Fusion Lumos (Thermo Fisher Scientific, Waltham, MA, USA). Survey scans were performed at R = 240,000 with wide isolation analysis of 300–1,350 mz. Data dependent top speed (1 second) MS/MS sampling of peptide precursors was enabled with dynamic exclusion set to 20 seconds on precursors with charge states 2 to 4. MS/MS sampling was performed with 1.6 Da quadrupole isolation, fragmentation by HCD with NCE of 25, analysis in the ion trap with maximum injection time of 10 milliseconds, and AGC target set to 3 x 10^4^.

#### Analysis

Raw files were analyzed using MaxQuant 1.6.0.1 (33, 34). Spectra were searched using the Andromeda search engine against a *Z. mobilis subsp. mobilis* ZM4 (GEO accessions CP023715, CP023716, CP023717, CP023718, CP023719) protein database and a target decoy database generated in house. Label free quantitation (LFQ) (35) and match between runs were toggled on, and ion trap tolerance was set to 0.4 Da. All other parameters were set by default. Peptides were grouped into subsumable protein groups and filtered to 1% FDR, based on target decoy approach. Downstream analysis of protein group LFQ values were performed using the Perseus software platform (36). First, all LFQ values were log2 transformed and any protein groups missing a value from ≥ 6 samples were removed followed by missing value imputation. Fold changes for each protein were calculated for the cellobiose and sucrose conditions by comparison of the LFQ against that of the control sample from the appropriate condition (intra- or extracellular). A Student’s two sample t-test was performed for each fold change measurement and p-values were corrected for multiple hypothesis testing by the Benjamini-Hochberg method to generate quantitative FDR values.

K-means clustering was performed on glucose normalized log2-fold change values for proteins that were measured in both the intracellular and extracellular fraction experiments (1199 proteins). The desired number of clusters was set to 50 using Euclidian distance and average linkage. Gene ontology enrichment in each cluster was performed using Fishers exact test with Benjamini-Hochberg correction for multiple hypotheses (p<0.05). Gene ontology annotations were downloaded from UniProt (37).

## SUPPORTING INFORMATION CAPTIONS

**S1 Table. Primers used in this study.**

**S2 Table. Plasmids and strains used in this study.**

**S3 Table. Localization prediction of glycosyl hydrolase used in this study.**

**S4 Table. Enriched gene ontology (GO) terms in both upregulated and downregulated intracellular and extracellular fractions of cellobiose- and sucrose-adapted *Z. mobilis* ZM4+pGH3 strain relative to glucose grown cells.**

**S1 Figure. Growth of *Z. mobilis* with GH3 and pVector control in rich medium glucose (RMG) supplemented with required antibiotics and IPTG.**

**S2 Figure. Growth of *Z. mobilis* GH3 on cellobiose requires IPTG.** The left plate (A) is RMC with 100 µg spectinomycin/ml and no IPTG. The right plate (B) is RMC with 100 µg spectinomycin/ml and 0.4 mM IPTG.

**S3 Figure. Heterologous protein production measurement.** (A) SDS-PAGE showing total crude proteins. (B) Highlighted showing GH3 produced in the sample with IPTG induction. (C) GH3 signal measured as percent of total signal in each lane.

**S4 Figure**. **Cellobiose conversion and ethanol production by *Z. mobilis* GH3 or pVector control in RMCG medium.** (A) Cellobiose conversion by *Z. mobilis* ZM4 strains containing pVector or pGH3 plasmids in RMCG medium. Cellobiose conversion was observed after 96 hours only in the strain expressing glycosyl hydrolase (pGH3) but not in the control with pVector. (B) Ethanol production was observed after 96 hours only in the strain expressing glycosyl hydrolase (pGH3) but not in the control with pVector. Some ethanol may have evaporated with escaping CO_2_ or during sampling for apparent OD_600_ measurement. Error bars are standard deviations of triplicate experiments.

**S5 Figure**. **Activity of cellular fractions of unadapted and adapted *Z. mobilis* GH3.** GH activity by different cellular fractions of the unadapted strain grown in RMG (A) and rich medium sucrose (B). Similarly, GH activity of different cellular fractions of cellobiose-adapted strain grown in RMCG (C), and GH activity of different cellular fractions of sucrose-adapted strain grown in RMCG (D). Abbreviations: Sup – supernatant, Peri – periplasmic fraction, Cyto cytoplasmic fraction, Sph – spheroplast and Wcells – whole cells (cell pellets).

**S6 Figure. Cellobiose conversion (A), ethanol production (B) and sucrose metabolism (C) by unadapted *Z. mobilis* GH3, pVector control, and adapted *Z. mobilis* GH3 in an RMC medium with increasing concentrations of sucrose (0.2-0.8%).** *Z. mobilis* containing pVector and pGH3 (pVect-G and pGH3-G, respectively) were pregrown in RMG medium. Similarly, *Z. mobilis* containing pVector and pGH3 (pVect-S and pGH3-S, respectively) were pregrown in RMS medium or increasing concentrations of sucrose (0.2%S – 0.8%S) in RMC. Samples were assayed at 0, 6, 24, 48, and 72 hours (legend on right).

**S7 Figure. Extracellular and intracellular proteomics of *Z. mobilis* GH3 grown in cellobiose and sucrose medium.** (A) Scatter plot of extracellular proteins of adapted strains grown on cellobiose or sucrose versus glucose, and (B) scatter plot of intracellular proteins of adapted strains grown on cellobiose or sucrose versus glucose. Proteins of interest are highlighted. Black proteins related to secretion and transport, Orange – glycosyl hydrolase (CC_0968). (C) and (D) Venn diagrams showing overlap between changes in proteins for cells grown on cellobiose or sucrose versus glucose (adjusted *p*-value <0.001).

**S8 Figure. Change in GH3 (CC_0968) level in cellobiose- and sucrose-adapted *Z. mobilis* GH3.**

**S1 File. Quantitative proteomics data.**

